# Toward precision EEG: Assessing the reliability of individual-level ERPs across EEG Systems

**DOI:** 10.1101/2025.11.10.687573

**Authors:** Adi Korisky, Eshsed Rabinovitch, Paz Har-Shai Yahav, Bruce D McCandliss, Elana Zion-Golumbic

## Abstract

Event-related potentials (ERPs) are among the most established tools for studying the neural mechanisms of perception and cognition. Advancing toward precision EEG, however, places new demands for a better understanding of how reliable neural markers are at the individual subject level. We conducted two complementary experiments to examine the reliability of N100 and P300 components in an auditory oddball paradigm with three sounds (Standard, Target, and Novel). In *Experiment 1*, we evaluated the consistency at both the group level and the individual level across four EEG systems: one research-grade wired system (BioSemi) and three mobile devices—Smarting, DSI-24, and EPOC X. At the group level, all systems demonstrated the canonical N100 and P300 components; however, the EPOC X system showed a significantly reduced signal-to-noise ratio compared to the others. At the individual level, temporal and spatial clustering analyses showed that N100 and P300 components were detectable in most individuals (70–85%), with additional significant responses appearing outside this range. We further calculated the similarity of individual responses across participants (“typicality index”), which revealed highly consistent responses to Standard and Novel sounds, alongside divergent patterns of responses to Targets. In *Experiment 2*, we assessed the within-participant reliability of N100 and P300 using a test–retest design. Results indicated high within-participant consistency of response patterns for all three stimuli, demonstrating that individual ERPs remain reliably stable over time, even when they deviate from canonical group-level patterns. The current study contributes to the ongoing discussion regarding the utility and reliability of ERP-based metrics for precision imaging and highlights important methodological considerations for their practical implementation.

## Introduction

Event-related potentials (ERPs) have long been a cornerstone of cognitive neuroscience, offering a robust, non-invasive, and cost-effective means of examining the neural dynamics that underlie perception and higher-order cognitive functions. Since its inception (Donchin, E. et al., 1973; Hillyard et al., 1973, 1998; Picton et al., 1974; Squires et al., 1975; Sutton et al., 1965), ERP research has relied heavily on averaging responses across individuals as a way to increase the signal-to-noise and to obtain neural responses that were common and generalizable (Luck, 2014; Makeig et al., 2004; Sur & Sinha, 2009). This approach has been hugely successful, leading to some of the most foundational discoveries in cognitive neuroscience and establishing a taxonomy of canonical ‘ERP components’ that have been used for decades to study the mechanisms underlying perceptual and cognitive processes in the human brain (Duncan et al., 2009; Hajcak et al., 2019; Helfrich & Knight, 2019; Luck, 2012, 2014). And yet, relying solely on group averages has its limitations, particularly if we strive to use ERP-based metrics to learn something about individual brains and their idiosyncrasies. This scientific aspiration, sometimes referred to as *precision imaging* framework, captures the desire to use neuroimaging tools for the purpose of individualized assessments of neurocognitive abilities or clinical conditions, with the capacity to inform personalized interventions and to assess treatment efficacy (Dubois & Adolphs, 2016; Gordon et al., 2017). Indeed, ERP-based measures hold much promise to serve as potential biomarkers for different neurocognitive conditions (de Aguiar Neto & Rosa, 2019; Keizer, 2021; McLoughlin et al., 2014; O’Sullivan et al., 2006), such as schizophrenia (Johnstone et al., 2013; Ren et al., 2021; Rosburg, 2018; Rosburg et al., 2008; Shen et al., 2020), attentional deficit disorder (Gamma & Kara, 2016; Hasler et al., 2016; Kaiser et al., 2020; Slater et al., 2022), post-traumatic stress disorder (Javanbakht et al., 2011; Johnson et al., 2013; Lewine et al., 2002), and anxiety (Al-Ezzi et al., 2020; Howe et al., 2014; Lars Thoma et al., 2020; Turan et al., 2002). The applicational appeal of individual ERP-based biomarkers is enhanced by the impressive advances of mobile EEG technology in recent years, which increases accessibility to affordable neural recording, bringing devices directly to the individual – in their home, hospital, clinic or school (Davidesco et al., 2023; Davidesco, et al., 2021; Gillis et al., 2022; Hölle et al., 2021; Hölle & Bleichner, 2023; Mathewson et al., 2024; Sabio et al., 2024; Xu et al., 2022).

However, the utility of individual-level ERPs as biomarkers is complicated by the inherent variability of these responses. As described extensively over the years, the spatiotemporal properties of individual-level ERPs are strongly influenced by multiple factors, including anatomical differences, electrode positioning, and head size (Luck, 2014; Makeig et al., 2004; McCarthy & Wood, 1985; O’Connor et al., 1994). Moreover, the method for identifying specific ERP components in individual responses can be challenging, often relying on visual peak-picking or general heuristics that do not always capture the extent of individual variability and lack standardized procedures (Davidesco et al., 2023). Adding to that, the various ERP components (e.g., N100, MMN, P2, P300, N400, P600, etc.), may differ in their detectability across individuals and specific design features. For example, the P300 components, which is often evoked by surprising events, is deemed more reliable than other components such as N100 or P2 (Fabiani et al., 1998; Intriligator & Polich, 1995; Polich, 1987, 1997; Segalowitz & Barnes, 1993; Sklare & Lynn, 1984; Tervaniemi et al., 1999). Hence, the assumption that ERPs of individuals look like ‘noisy’ versions of group-level ERPs, in terms of time-course and spatial topography, does not seem to hold and poses a stark barrier for ERP-based precision imaging (Clayson et al., 2019; Fields & Kuperberg, 2020; Hajcak et al., 2017; Höller et al., 2017; Jensen & MacDonald, 2023; Karvelis et al., 2023; Luck & Gaspelin, 2017; Melnik et al., 2017). Specifically, it makes it difficult to distinguish between individual differences that truly reflect variability in neurocognitive abilities or clinical states (and accordingly could potentially serve as meaningful biomarkers) vs. individual differences that simply capture the “natural” variability between healthy brains (Hajcak et al., 2017; Höller et al., 2017). That said, if we had a good assessment of this “natural” variability in individual-level ERP components, this would substantially advance efforts to assess their utility for precision imaging. The current study is an attempt to do just that – to quantify the similarity and variability in ERP responses across individuals, for specific ERP components and for the entire spatio-temporal pattern (Segalowitz & Barnes, 1993; Tomé et al., 2015).

We focused on one of the most well-studied ERP paradigms - the 3-sound auditory oddball task (Fabiani et al., 1987; Hillyard et al., 1973; O’Connor et al., 1994; Squires et al., 1976). In this paradigm, participants listen to a sequence of repeated tones (Standards) and are asked to detect specific deviant tones (Targets). In addition, occasional Novel sounds are presented (in this case, short ecological sounds such as phone ringtones; Masson and Bidet-Caulet 2019), which are not expected. This paradigm yields two prominent ERP components – an early N100, that reflects the neural response in early auditory cortices (Coull, 1998; Hillyard et al., 1973, 1998; Oatman & Anderson, 1977) and a later P300 response, that is associated with higher cognitive processes including attention-capture, stimulus discrimination and decision-making (Coull, 1998; Linden, 2005; Ghani et al., 2020; Hillyard et al., 1973; Lee et al., 2014; Polich, 2007; Tomé et al., 2015). While the N100 is considered an obligatory response for all auditory stimuli, the P300 is more selective and is usually observed in response to deviant, target, or surprising events (with potential dissociations between P300 subcomponents. For review, see Polich, 2007). Here, we use this paradigm to assess the similarity and variability in ERP responses to Standard, Target, and Novel sounds, in a non-clinical population, focusing on the N100 and P300 components as well as on the ERP waveform.

We conducted two complementary experiments. In **Experiment 1**, we assessed the consistency/variability of the time-course of neural responses across individuals and the probability of identifying the N100 and P300 components at the individual-level, based on group-constrained or fully data-driven analyses. Specifically, we compared results across *four different EEG systems* – spanning wired and wireless systems, research-grade and consumer-grade, with different numbers of sensors. These included: BioSemi (64-channel, wired, gel-based); Smarting (24-channels, wireless, semi-dry by mBrainTrain); DSI-24 (24-channels, wireless, dry, by Wearable Sensing), and EPOC X (14-channels, wireless, semi-dry, by Emotiv). By conducting the same experiment using different EEG systems and in different experimental setups, we aimed to assess the generalizability of individual-level ERPs, and their robustness across contexts and technical specifications of different systems. This effort aligns with growing efforts to advance the use of mobile neurotechnologies for clinical and research purposes outside the lab (For comprehensive reviews see: Larsen et al., 2024; D’Angiulli et al., 2022; Hölle et al., 2021; Janssen et al., 2021; Lau-Zhu et al., 2019; Mathewson et al., 2024; Niso et al., 2023; Xu & Zhong, 2018). **Experiment 2** was conducted using a research-grade EEG system (BioSemi Active II) and was aimed at evaluating the degree of within-subject ERP consistency and how it related to between-subject variability. Participants repeated the same oddball task experiment as in Experiment 1 twice, in a test-retest design, and we quantified the similarity of individual ERP spatio-temporal morphology between runs.

## Methods

### Experiment 1

#### Participants

EEG was recorded from 37 participants (23 females, 16 males), aged between 15 and 32 (see Table 1 for description of the sample tested with each EEG system). Most participants were right-handed (4 with a dominant left hand) and reported having no hearing problems or diagnosis of any neurological or psychiatric condition. The study was approved by the Institutional Review Board (IRB) of Bar-Ilan University. Informed written consent was obtained from all participants. For participants under the age of 18 (see the EPOC X setup), additional ethical approval was granted by the Israeli Ministry of Education, and parental consent was required before each session. All participants were compensated for their participation with money, course credits, or small tokens.

**Table 1:**
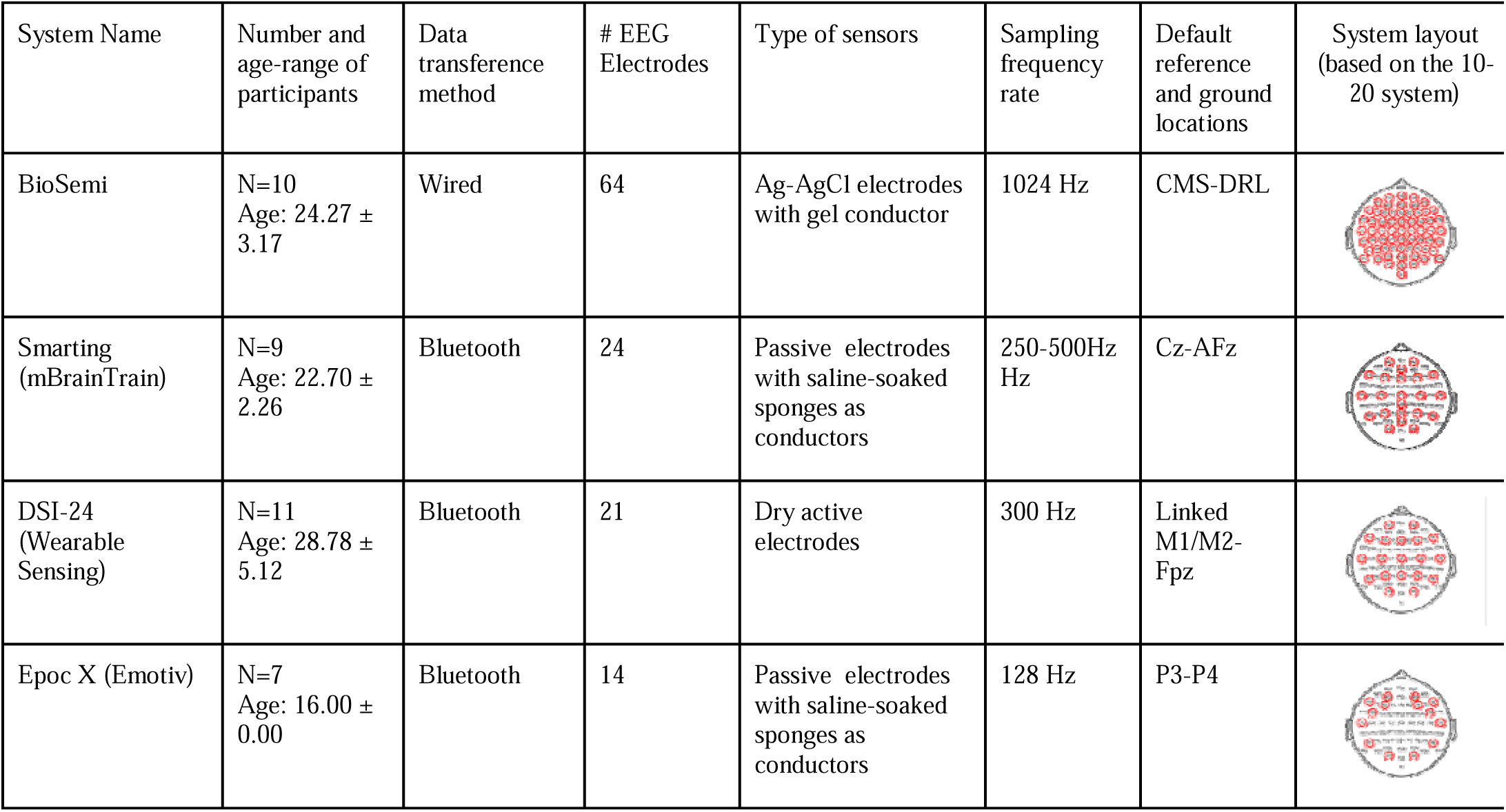
Overview of the four EEG systems used in the study. The table presents participant demographics for each setup and summarizes key characteristics of each system, including data transfer method, number and type of electrodes, sampling frequency, and default reference and ground locations. Scalp layouts in the right column illustrate the electrode positions for each system (circled in red), overlayed on the scalp layout of the BioSemi channel system (based on the international 10–20 system).

| System Name | Number and age-range of participants | Data transference method | # EEG Electrodes | Type of sensors | Sampling frequency rate | Default reference and ground locations | System layout (based on the 10-20 system) |
| --- | --- | --- | --- | --- | --- | --- | --- |
| BioSemi | N=10<br>Age: 24.27 ± 3.17 | Wired | 64 | Ag-AgCl electrodes with gel conductor | 1024 Hz | CMS-DRL | |
| Smarting (mBrainTrain) | N=9<br>Age: 22.70 ± 2.26 | Bluetooth | 24 | Passive electrodes with saline-soaked sponges as conductors | 250-500Hz Hz | Cz-AFz | |
| DSI-24 (Wearable Sensing) | N=11<br>Age: 28.78 ± 5.12 | Bluetooth | 21 | Dry active electrodes | 300 Hz | Linked M1/M2-Fpz | |
| Epoc X (Emotiv) | N=7<br>Age: 16.00 ± 0.00 | Bluetooth | 14 | Passive electrodes with saline-soaked sponges as conductors | 128 Hz | P3-P4 | |

#### Paradigm

All participants performed an auditory oddball task, designed as follows. Stimuli consisted of three types of auditory stimuli: Standard tones (1000 Hz, 50 milliseconds duration), Target tones (1500 Hz, 50 milliseconds duration), and a variety of Novel ecological sounds (300 milliseconds duration), such as phone ringtones, adapted from the work of Masson and Bidet-Caulet (2019). All stimuli included 10-millisecond ramp-up and ramp-down phases.

Figure 1 illustrates the structure of the task. In each block, 80 auditory stimuli were presented, which included 70% Standard tones, 15% Target tones, and 15% Novel sounds. Participants were instructed to respond via keyboard button press only to the Target tone. Stimulus order was pseudo-randomized with two constraints: (1) at least five Standard tones were presented consecutively at the beginning and the end of each block; (2) at least one Standard tone was interposed between a Target tone and a Novel sound. We used a constant inter-stimulus interval (ISI) of 1000 milliseconds, and each block lasted ∼87 seconds. The experiment contained one initial training block followed by eight experimental blocks, yielding a total of ∼450 Standard tones, 96 Target tones, and 96 Novel Sounds. A short break was given between each experimental block, and the onset of the next block was self-paced. The experiment was programmed and presented using the PsychoPy software platform (Peirce, 2007).

**Figure 1:**
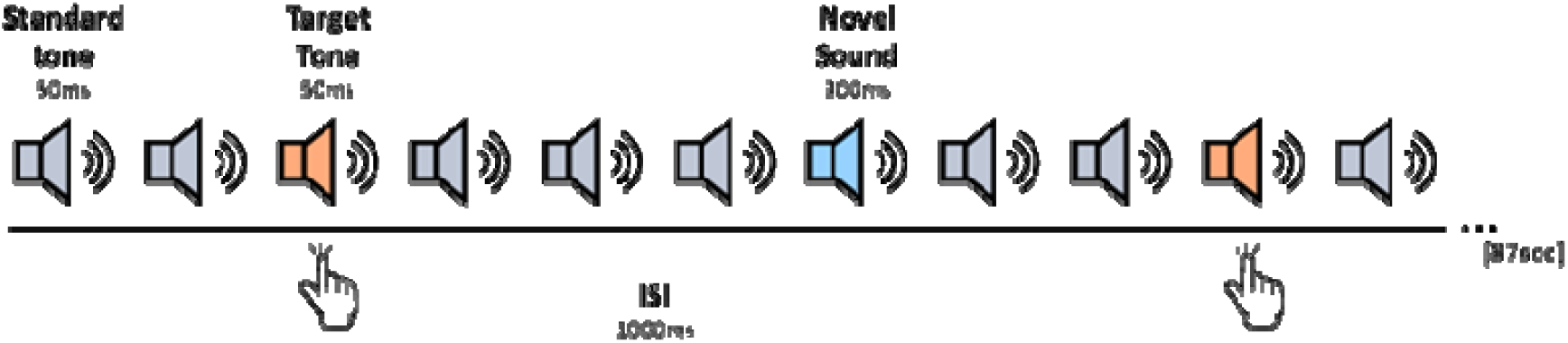
Experimental design of the Auditory Oddball paradigm. Each block consists of 80 auditory stimuli in the following division: 70% Standard tones (1000 Hz), 15% Target tones (1500 Hz), and 15% Novel ecological sounds (e.g., ringtones). Stimuli were presented with a fixed 1000 ms ISI. The experiment included one training block and eight experimental blocks.

#### EEG recording (per system)

EEG data were recorded from four different systems (N=7-11 participants per system, see Table 1 for a detailed description). The same oddball paradigm was used for all systems; however, some aspects of the experimental setup varied from system to system, as a function of their technical specifications as well as other experimental constraints. Rather than viewing these variations as an obstacle for comparison, we see them as a crucial component of our overall scientific goal, which is to assess the generalizability of ERP-based markers across contexts and EEG systems. Similarly, our choice to test a relatively small sample with each system aligns with our overarching goal to evaluate the sensitivity for identifying replicable neural response within-subject. Our statistical analyses focus primarily on inter-subject statistics, focusing on the similarity between participants rather than relying on group-level averaging. Below, we provide details of the specific experimental setup used for each system.

##### BiosSemi

We used the BioSemi Active II EEG system (BioSemi BV, Amsterdam, Netherlands; https://www.biosemi.com/Products_ActiveTwo.htm) as our ‘gold standard’ to which three mobile systems were compared. We used a gel-based 64-channel system, with Ag-AgCl electrodes positioned according to the 10-20 system and a 1024 Hz sampling rate. Data was recorded in an electrical-shielded and acoustically-attenuated room in a lab at Bar Ilan University. Participants were seated comfortably in front of a computer screen, where visual instructions and a central fixation cross were presented. Auditory stimuli were presented in a free-field manner through a loudspeaker positioned in front of the participant. EEG data were recorded through the Lab Streaming Layer platform (LSL) (Kothe, Christian et al., 2014). The audio in the room during the experiment was recorded using an external microphone and also streamed into LSL (via the audio-capture interface; https://github.com/labstreaminglayer/App-AudioCapture) to facilitate accurate segmentation of the EEG data based on the actual audio perceived.

##### Smarting

The second EEG system tested was a saline-based wireless Smarting system (mBrainTrain LLC, Belgrade, Serbia; https://mbraintrain.com/), with 24 EEG electrodes positioned according to the 10-20 system and a 250 Hz sampling rate. Data was recorded using a similar setup as the BioSemi data, in the same electrical-shielded and acoustically-attenuated room at Bar Ilan University. EEG data was wirelessly streamed to the recording computer via Bluetooth and was recorded using LSL, and synchronized to the audio recordings from the microphone (described above).

##### DSI

The third EEG system tested was the Wearable Sensing DSI wireless dry-electrode system (Wearable Sensing, San Diego, CA, USA; https://wearablesensing.com/dsi-24/), featuring 21 active sensors, positioned according to the 10-20 system and using a 300 Hz sampling rate. These data were collected in a field-based setting, in a quiet yet non-shielded classroom at Stanford University, California, USA. Here, participants completed the experimental task on a laptop, and auditory stimuli were delivered in a free-field manner through the laptop’s built-in speakers. The audio in the room during the experiment was recorded using the internal microphone on the laptop. EEG data was wirelessly streamed to the recording computer via bluetooth and was recorded using LSL, and synchronized to the audio recordings from the microphone.

##### EPOC X

The last EEG system tested was the EPOC X system by Emotiv (Emotiv Inc., San Francisco, CA, USA; https://www.emotiv.com/), which features 14 passive saline-based electrodes, positioned according to the 10-20 system and using a 128 Hz sampling rate. These data were collected in a field-based setting, in a quiet yet non-shielded classroom in a local high school (in-school lab). Participants were 9th-grade students, aged 14–15, and they completed the experiment as part of an ongoing neuroeducation research-practice partnership between the Begin High School in Ramat Gan and the research team from Bar Ilan University (Korisky et al., 2024). In this setup, participants performed the task on a laptop and auditory stimuli were delivered through in-ear headphones (to avoid excessive noise from outside sources). EEG signals were wirelessly streamed to the recording computer via Bluetooth, recorded using LSL, and synchronized to the audio recordings from the microphone.

### Data Analysis

#### Behavior Analysis

Key presses were analyzed to classify responses as hits, misses, or false alarms as follows: a key press was classified as a ‘hit’ if it occurred within 200 to 1500 milliseconds after the target tone onset. Otherwise, that target was considered to be ‘missed’. All other key presses were considered ‘false alarms’, indicating responses made mistakenly or unrelated to the stimuli.

#### EEG Analysis

Data analysis protocols were highly similar across all EEG systems, and were based on the MATLAB-based FieldTrip toolbox (MathWorks 2021, https://www.mathworks.com; Oostenveld et al., 2011) using identical scripts, with only slight adaptations due to system-specific differences (e.g., file format, layout etc.).

#### Preprocessing

Raw EEG data were re-referenced to linked right and left mastoids (BioSemi, Smarting and EPOC X systems) or to linked left and right ear lobes (DSI system, default). Then, data were bandpass filtered between 0.5 and 40 Hz and detrended and demeaned to retain the frequency range associated with auditory ERPs. For artifact correction, we employed independent component analysis (ICA) to remove ocular and cardiac artifacts (identified through visual inspection of the time course and spatial distribution of the ICA components).

The onset of each auditory stimulus was identified using combined indications from digital triggers sent from PsychoPy and the audio recordings. The continuous EEG data were epoched into trials ranging from −100 to 700 ms around the onset of each sound. Epochs with remaining muscle-related or other artifacts or high noise were identified and rejected based on the standard deviation of the EEG signal (STD) using the ft_rejectvisual function in the FieldTrip toolbox (thresholds for rejection were determined separately for each EEG system: BioSemi and Smarting - 25 STD; DSI - 35 STD; EPOC X - 40 STD). The number of trials rejected from each system in each condition is reported in Table 2.

**Table 2:** The number of trials rejected for each condition and EEG recording system (mean ± STD).

| Condition | BioSemi | Smarting | DSI | EPOC X |
| --- | --- | --- | --- | --- |
| Standard | 22.2 ±12.60 | 39.3 ±27.01 | 12.3 ±11.57 | 40.8 ±27.41 |
| Target | 3.7 ± 4.34 | 6.4 ±5.66 | 1.3 ±2.11 | 7.1 ±5.06 |
| Novel | 5.1 ±5.19 | 6.8 ±6.55 | 1.2 ±1.23 | 8.5 ±6.95 |
| Mean across conditions | 10.4±10.31 | 17.5±18.88 | 4.9±6.40 | 18.8±19.10 |

##### Group Level Analysis

Clean epochs were averaged for each participant, separately for Standard, Target, and Novel stimuli. Averages were then low-pass filtered at 12 Hz (4th order zero-phase Butterworth filter), and baseline-corrected to the pre-stimulus period (−100 to 0 ms for BioSemi, Smarting and DSI-24 and 50 – 150 ms for EPOC X, following a visible delay in data acquisition) to produce ERPs. Grand average responses were derived by averaging the ERPs across participants, separately for each system and stimulus type.

To identify time windows where the ERP significantly deviated from zero, we used a data-driven clustering approach for each system. For each task condition, a one-sample t-test was performed at each time point to identify periods where the response differed significantly from zero (p < 0.05). Next, a temporal clustering permutation test was applied to identify contiguous time windows with significant responses at each electrode (Maris & Oostenveld, 2007). In addition, we visually inspected the grand-average waveforms to identify peaks that represent the N100 and P300 components based on their timing, polarity, and scalp topography.

To assess the similarity of ERP responses across systems, we calculated the cross-correlation between the grand-average ERP waveforms from each mobile EEG system and those obtained with the BioSemi cap, which served as the lab-standard reference. This analysis was restricted to ERPs from the one centro-frontal electrode (‘F4’ for EPOC X and ‘Fz’ for all other systems) where the N100 and P300 components were maximal. For each between-system comparison, we extracted the temporal lag corresponding to the maximum correlation and the associated correlation coefficient (Pearson’s r) at this lag.

##### Individual Level Analysis

The individual-level approach aimed to examine whether the N100 and P300 components that were identified at the group level could also be reliably detected within individual participants’ ERPs. Given the inherent variability in the spatio-temporal morphology of neural responses across individuals, we employed three complementary within-subject analyses that vary in the a-priori assumptions that they impose from the group-level analysis onto the individual-level data. These analyses were applied to single-trial data from each condition and are reported separately for each EEG system.

The **first** analysis sought to determine the consistency of the time-course of neural responses to each stimulus across individuals, beyond spatial variations between them, and their relation to the group-average. To this end, single-trial EEG data from each participant, one-tailed t-tests were performed at each electrode and each time-point (binned into 20-ms windows to reduce multiple comparisons). Using a statistical threshold of p < .05 (uncorrected), we assessed how many participants showed significant responses at any given point in time, across all electrodes and at specific electrodes. We specifically quantified the proportion of participants showing significant neural responses within the time-windows identified in the group-averages for the N100 and P300 peaks.

The **second** analysis used a more rigorous statistical analysis to identify the N100 and P300 peaks in individual ERP time courses, and to assess the similarity of the time-course of neural responses across individuals. This analysis was restricted to a single electrode where these responses were maximal in the group-averages (Fz or the closest available electrode), in order to reduce the number of multiple comparisons and to overcome potential bias due to the uneven number of electrodes across EEG systems. For each participant, we averaged the single-trial EEG data in each condition to obtain their personal ERP. We applied a two-way t-test to the single-trials at each time-point and used a temporal clustering permutation test (permutest.m function, MATLAB, Gerber (2025), https://www.mathworks.com/matlabcentral/fileexchange/71737-permutest) to identify clusters of consecutive time points where the ERP differed significantly from zero (positive or negative, alpha < 0.025). This statistical approach captures the truly individuated time-courses of neural responses for each participant. Next, we examined the consistency of ERP time courses across individuals by calculating the pairwise correlations between the ERPs of all participants (recorded with the same EEG system), and computed a “typicality score” for each participant: the average of their correlations with all other participants. We further tested whether this measure varied across conditions and EEG systems using a mixed repeated-measures 2×3 ANOVA, with condition (standard, target, novel) as a within-subjects factor and system (BioSemi, Smarting, DSI-24, EPOC X) as a between-subjects factor. Bonferroni-corrected post hoc comparisons were used to assess pairwise differences.

The **third** analysis focused on the similarity in the spatial distribution of neural responses across individuals. This analysis was restricted to the two time-windows identified in the group-level analysis for the N100 and P300 components (See Table 3) to reduce the number of multiple comparisons. For each participant, we averaged the single-trial EEG data within these time-windows at each electrode and applied a one-way t-test at each electrode with a spatial-clustering permutation test to identify clusters of consecutive time points where the response differed significantly from zero (negative for N100, positive for P300; alpha < 0.025). Complementing the previous analysis, the approach allows us to investigate the extent to which individual topographies aligned with the group-level patterns.

**Table 3:** Time windows (in milliseconds) showing significant clusters (p < .05, cluster-corrected) corresponding to the N100 and P300 components in response to each task condition, presented separately for each system. Values in parentheses indicate the peak latency identified through visual inspection.

|  | N100 |  |  | P300 |
| --- | --- | --- | --- | --- |
|  | Standard | Target | Novel | Novel |
| BioSemi | 51 – 106 (77) | 50 – 100 (78) | 50 – 104 (87) | 233 – 327 (272) |
| Smarting | 40 – 104 (76) | 48 – 104 (80) | 32 – 168 (80) | 252 – 332 (296) |
| DSI-24 | (-16) – 43 (13) | (-6) – 50 (23) | (-13) – 66 (26) | 213 – 270 (240) |
| EPOC X | n.s. (277) | n.s. (269) | n.s. (269) | n.s. (497) |

Together, these analyses provide a basis for comparing and quantifying similarities and differences in ERP responses across participants, conditions, and EEG systems.

## Results

### Experiment 1

#### Behavior

Analysis of behavioral performance served as a means to confirm that participants were appropriately engaged in the task and to ensure similar behavioral characteristics across participants tested with different EEG systems. Due to a technical problem during the session, the behavioral data from the EPOC X system was unavailable for two participants, limiting the analysis to five participants (out of seven). The overall hit rate was relatively high (90% average across systems); and false alarms were minimal (average 2.53 false alarms per participant), indicating that participants tested with all systems understood the task and performed it well. Notably, performance was slightly lower for participants tested with the EPOC X, who were high school students tested in an in-school lab environment, as opposed to participants tested with the other systems who were older. Reaction times were longer for data collected using the DSI-24 and EPOC X systems (∼684 ms and ∼642 ms), both of which were collected under field conditions, relative to the BioSemi and Smarting systems (∼452 ms and ∼469 ms, respectively), which were collected in a lab setting. These differences tentatively point to potential effects of the testing environment on performance, a topic that should be taken into consideration, particularly when using mobile testing systems (see Figure 2).

**Figure 2:**
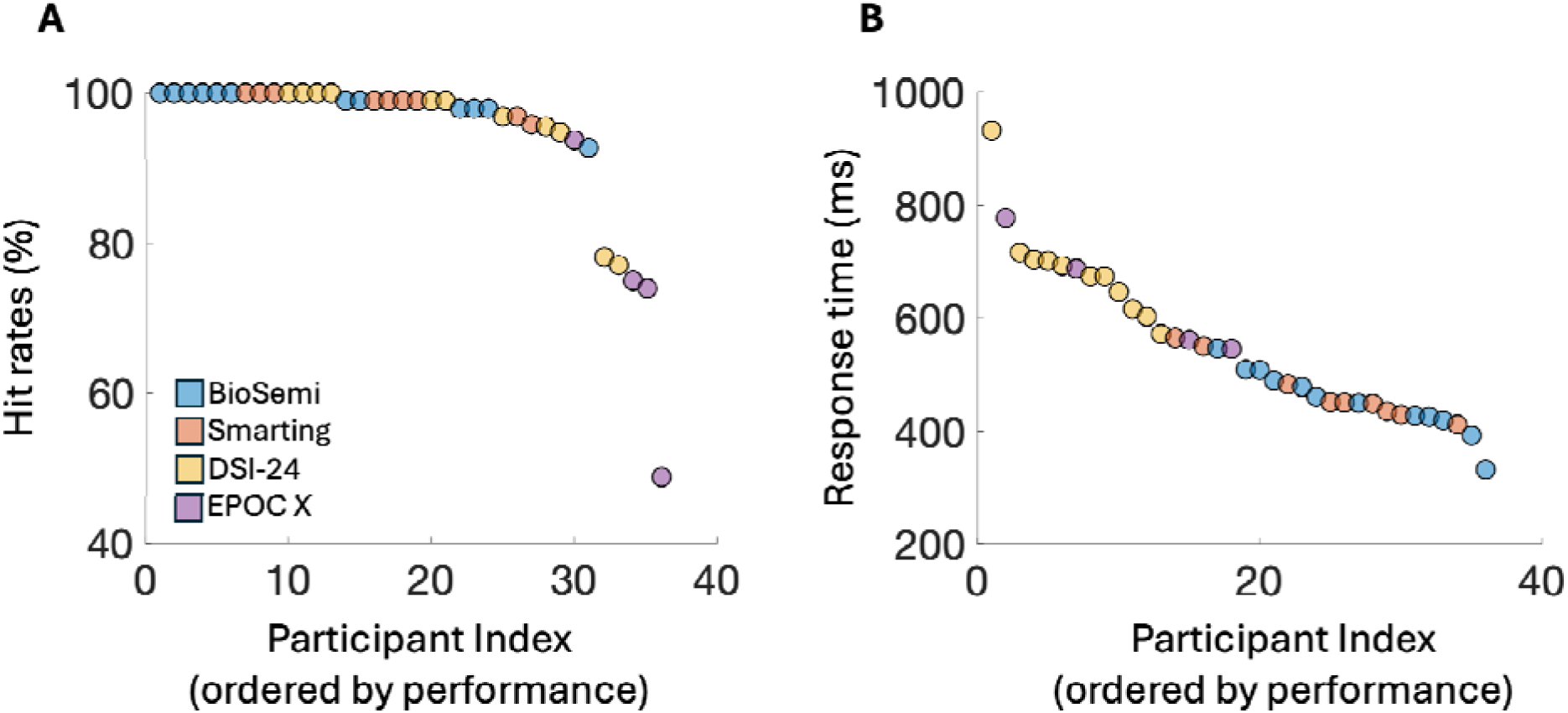
Behavioral performance on the Oddball tasks across EEG systems, showing (A) Mean hit rates, (B) Response time (ms). The X-axis represents the participant index, ordered by performance for each measurement, with each point corresponding to an individual participant.

#### EEG results

##### Group-level analysis

To identify group-level time windows for our targeted components – N100 and P300, we first explore the grand-average ERPs elicited in response to Standard, Target, and Novel sounds for each of our systems. Using cluster-based analysis, we identified clusters in the time course where the signal differed significantly from zero (at electrode ‘Fz’ or the closest one to it, p < 0.05, temporal-cluster corrected). Visual inspection reveals a replicable canonical shape of auditory ERPs in all four systems, showing an early frontocentral negative peak in response to all stimuli, corresponding to the N100 component, followed by a wide frontocentral positive peak only in response to Novel sounds, corresponding to the P300 component. Results also revealed differences in onset latencies across systems, with the EPOC X showing a pronounced delay. Since stimulus onset was estimated from audio recordings synchronized to the EEG data via LSL, this delay may be attributed to hardware limitations or data transmission latency.

**Figure 3:**
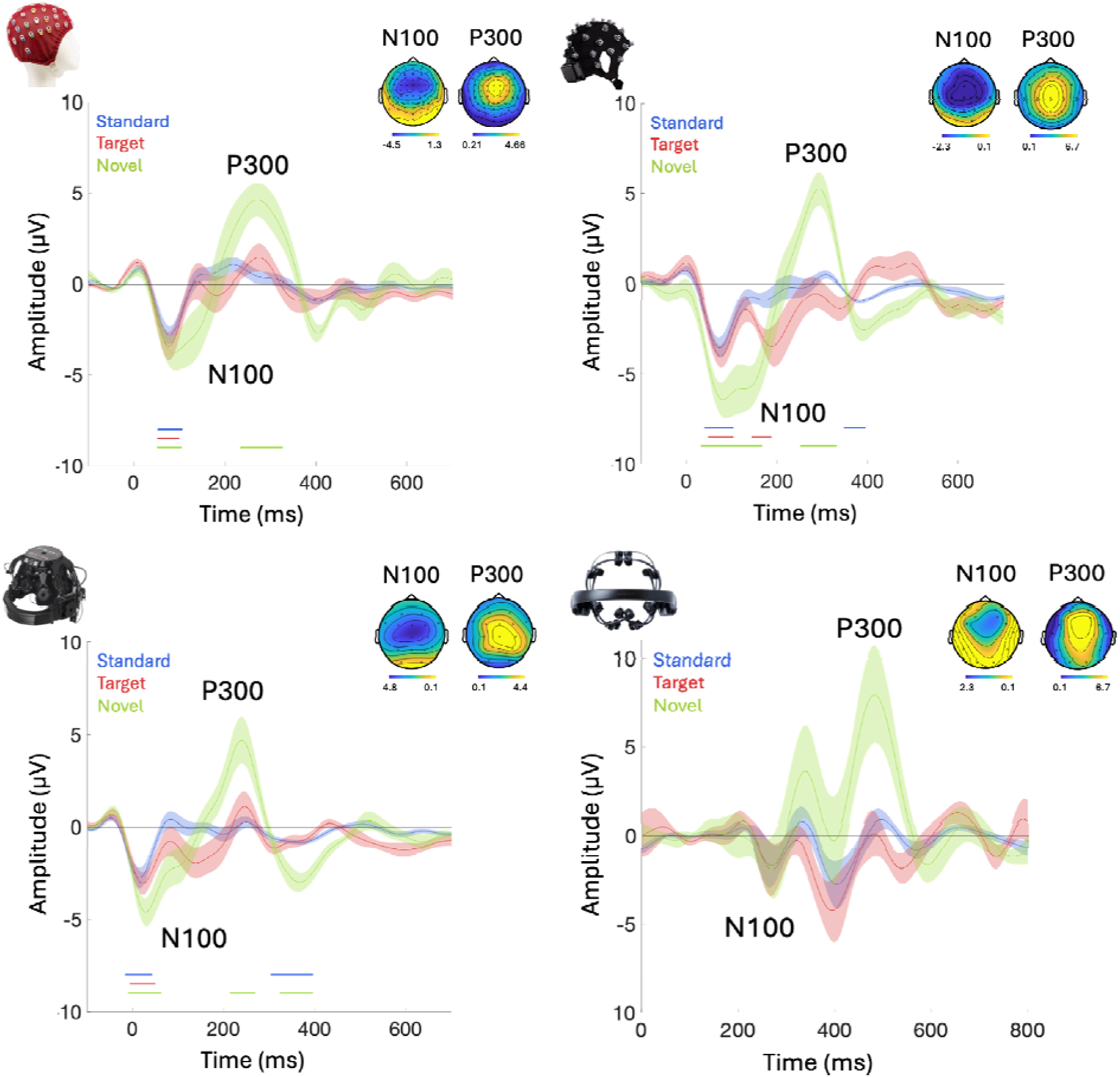
Grand-average responses and topographies of ERPs across systems: **(A)** BioSemi (n=10), **(B)** Smarting, mbt (n=9), **(C)** DSI-24, Wearable Sensing (n=11), **(D)** EPOC X, EMOTIV (n=8). Each panel presents the grand average ERP elicited in response to the three stimuli: Standard, Target, and Novel sounds, from one selected electrode (‘F4’ for EPOC X, ‘Fz’ for all other systems). Shaded areas around the curves represent the standard error across participants. Time t=0 represents our best estimation of the onset of the stimulus, which was determined based on direct audio-recordings that were synchronized to the EEG recordings via LSL (see Methods). Nonetheless, some timing differences are observed between the systems, most notably with greatly delayed responses with the EPOC X system. Thus, for the BioSemi, Smarting, and DSI, the baseline was computed on 100 ms before the onset of the stimuli while the baseline period for EPOC X was computed from 50 ms to 150 ms. The colored horizontal lines beneath each figure indicate the time-windows where the ERP in response to each stimulus differs significantly from zero (p < 0.05, cluster corrected). The topographical distribution of the N100 and P300 responses is presented in the inset of each panel (±10 ms around each peak; identified based on visual inspection). For visual comparison purposes, topographies are shown for all identified peaks, regardless of whether they reached statistical significance in the cluster analysis.

To assess the similarity between the ERPs extracted using the mobile EEG systems to the BioSemi lab-grade device, we calculated the cross-correlation between them (which accounts for any potential temporal lags between systems). In Table 4 we report the correlation coefficients and time-lags showing maximal correlation between each of the mobile systems and the BioSemi, separately for the Standard, Target and Novel stimuli. ERPs measured with the Smarting system had a minimal time-lag between them (0-4 ms), owing to the fact that they were recorded in the same experimental setup, whereas the DSI-24 system had a lag of ∼ -50ms relative to the BioSemi system, which was roughly consist for the three stimuli (the *negative* lag implies supposedly *delayed* detection of stimulus-onset for segmentation purposes). Conversely, the EPOC X had a most substantial and variable lag, estimated to be between 203 ms (Target) and 328 ms (Standard). The existence of these time-lags between different experimental setups suggests that although the EEG segmentation procedure was identical for all four systems, and relied on direct recordings of the audio synchronized to the EEG via LSL, this procedure is still not immune to hardware or Bluetooth-related delays that are unique to each system and can result in mismatched timing between the EEG and audio signals. These results emphasize the importance of evaluating and calibrating the temporal delays, particularly in wireless systems (Bleichner et al., 2016; Hairston et al., 2014; Hairston, 2012; Ries et al., 2014).

**Table 4:** Maximal cross-correlation values (Pearson’s r) between the ERP grand averages obtained with BioSemi vs. each of the mobile EEG systems. The time-lag where the maximal cross-correlation was found for each comparison is shown in parentheses.

|  | Standard | Target | Novel |
| --- | --- | --- | --- |
| BioSemi vs. Smarting | $r = 0.73$ (4 ms lag) | $r = 0.41$ (0 ms lag) | $r = 0.87$ (4 ms lag) |
| BioSemi vs. DSI-24 | $r = 0.84$ (-63 ms lag) | $r = 0.58$ (-47 ms lag) | $r = 0.88$ (-50 ms lag) |
| BioSemi vs. EPOC X | $r = 0.65$ (328 ms lag) | $r = 0.4$ (203 ms lag) | $r = 0.66$ (227 ms lag) |

After correcting for these temporal lags, we found that ERPs to both Standard and Novel sounds, the Smarting and DSI-24 systems were strongly correlated with ERPs from the BioSemi (Pearson’s r between 0.73-0.88, p < .01) whereas the EPOC X system showed weaker but still significant correlations (Pearson’s r = 0.65 for both stimuli, p < .05). Interestingly, in all three systems, the correlations with responses to the Target stimuli were substantially lower (Pearson’s r between 0.4-0.58 for all systems).

##### Individual-level analysis

Guided by the group-level results, we now turn to inspect the ERPs of individual participants in response to each stimulus, across the different EEG systems. Figure 4 shows the proportion of participants who had a significant response at each time-point (vs. zero; one-sided p < .05, uncorrected here to reduce false-negatives), shown for all electrodes (Figure 4a) and at electrode ‘Fz’ (or ‘F4’ for the EPOC-X) where ERPs are maximal (Figure 4b). Visualizing individual-level ERPs in this way highlights both the similarities and differences between them. On the one hand, and not surprisingly, the time-windows in which the highest proportion of participants showed significant non-zero responses were those identified in the group-level analysis for the N100 and P300 responses (indicated in horizontal lines in Figure 4b), a pattern that was similar across EEG systems (bearing in mind the time-lags between systems, identified in the group-level analysis). At the same time, even in those time windows, only between 70% and 85% of participants had significant responses, demonstrating the limited generalizability of using group-derived metrics for identifying specific responses in individual-level ERPs. Moreover, there were **<u>no</u>** time-points (other than the baseline period) in which **none** of the participants had a significant response in any of the channels, and several time-points not indicated in the group-level average where over 50% of participants showed significant responses (Figure 4b). Together, these results emphasize the vast underlying variability of individual-ERPs, which is not captured by group averages.

**Figure 4:**
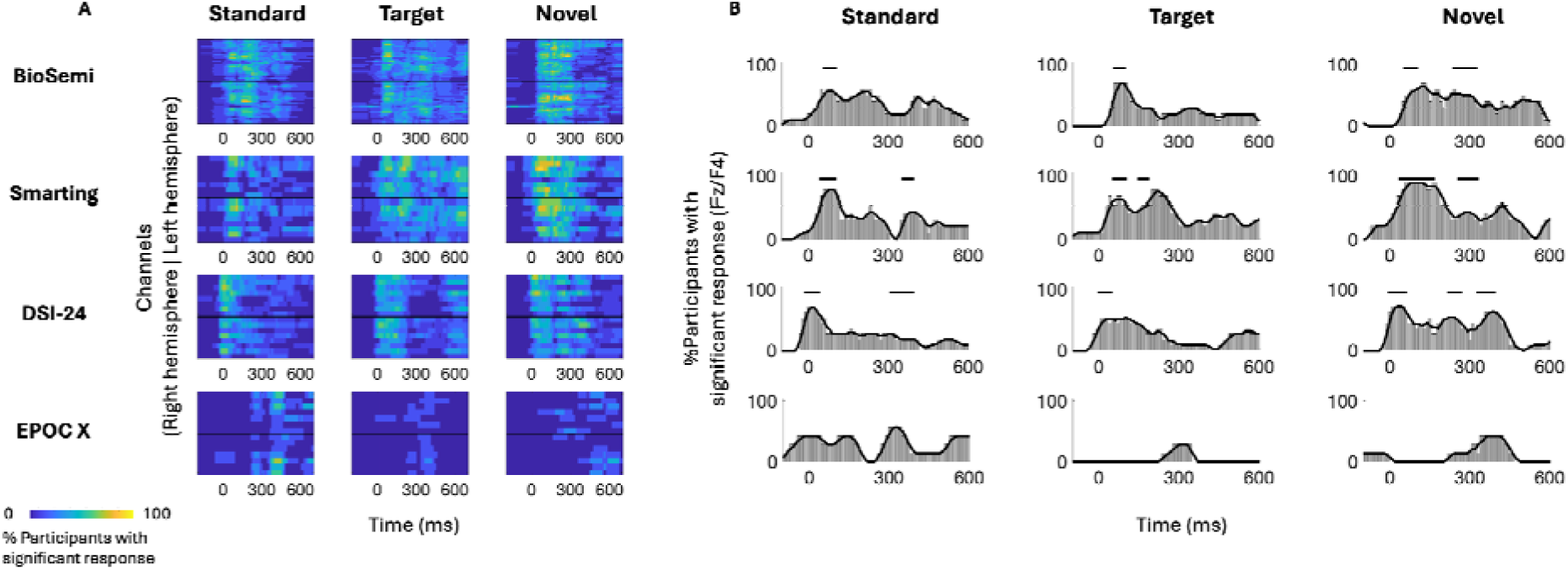
Whole-head analysis of individual ERP responses across participants. (A) Heatmap showing the percent of participants with a significant response at each time point and EEG channel (p < 0.05 t-test, uncorrected). The colors represent the percentage of participants (from 0% to 100%) that show a significant response in this window. The y-axis represents the channels of each system, organized from anterior (top) to posterior (bottom) in each hemisphere, with the black vertical line separating the right and left hemispheres. (B) Time course showing the percent of participants with a significant response from one chosen channel (‘F4’ - EPOC X and ‘Fz’ all other systems), to all stimuli. Horizontal lines represent the significant time windows that were identified in the group-level analysis (see Group-level analysis).

To further quantify this individual variability, we conducted more rigorous within-subject statistical analyses to identify the time-windows and spatial distribution of significant responses to Standard, Target, and Novel stimuli and their similarity across participants. Panel A in Figure 5-8 shows the ERPs from individual participants in response to each of the stimuli for each EEG system (from electrode Fz/F4 as appropriate), with horizontal lines indicating the time-windows identified as having a significant response (vs. zero; p < 0.05, temporal cluster corrected). Panel B in Figures 5-8 shows significant clusters of electrodes identified in the N100 time window (all stimuli) and P300 time window (Novel stimuli). These results are also summarized in Table 5, indicating that using this data-driven approach, significant N100 (all stimuli) and P300 response (Novel stimuli) could be identified in ∼50-80% of participants, after correcting for multiple corrections, with substantial variability across EEG systems. The likelihood of detecting significant ERPs in individual participants was better using the spatial-clustering approach (which was constrained to the time-windows identified in the group-level average) relative to the temporal-clustering approach (which was constrained to a single electrode with peak response in the group-level average). Comparison between the systems showed highest detection rates of individual-level ERPs in the BioSemi and Smarting data (70-100% and 89-100% respectively, using spatial clustering), and relatively good detection rates for DSI-24 data (72-81%), but quite poor results for the EPOC X data (14-42% using spatial clustering; 0-71% using temporal clustering). Taken together, this detailed perspective on individual-level ERPs suggests *a moderate level of sensitivity* for detecting these well-known responses in the EEG data of individuals, which is affected by the intrinsic spatio-temporal variability in the pattern of responses across individuals, as well as by the quality of the EEG signal itself.

**Table 5:**
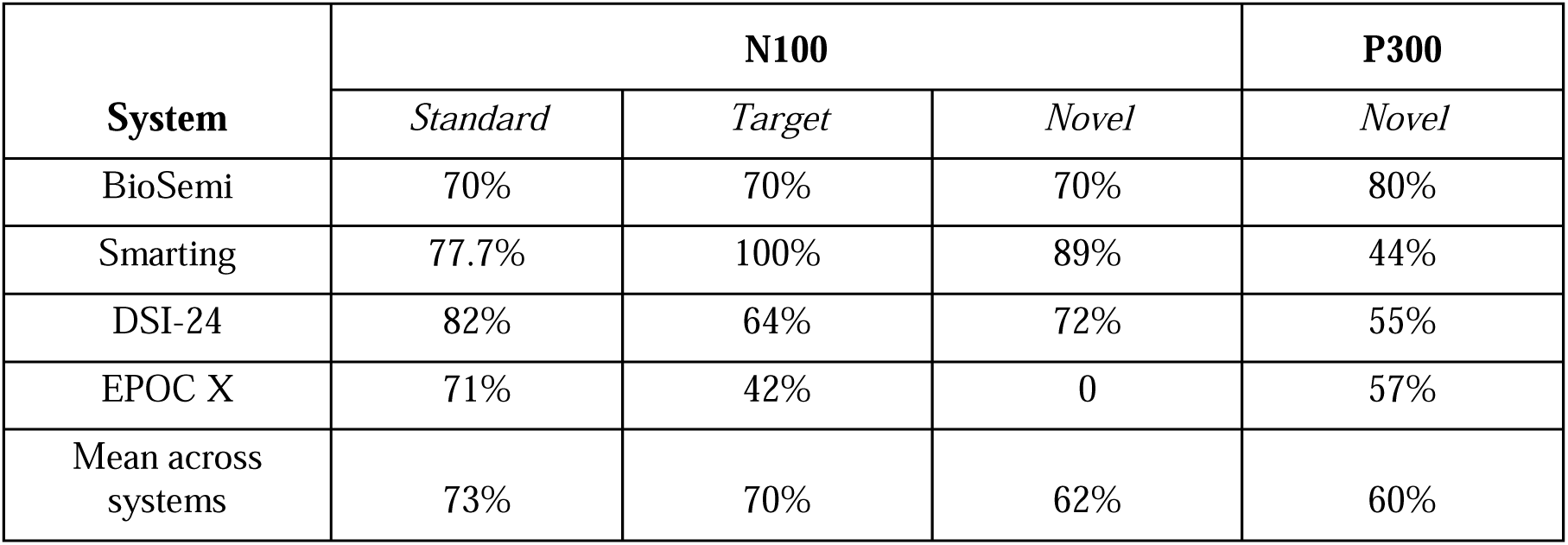
Temporal-clustering results. Percentage of participants exhibiting a significant cluster (p < 0.05 cluster corrected) from one chosen channel (‘F4’ - EPOC X and ‘Fz’ all other systems) corresponding to the N100 and P300 components in response to each stimulus, separately for each system.

| **System** | **N100** |  |  | **P300** |
| --- | --- | --- | --- | --- |
|  | *Standard* | *Target* | *Novel* | *Novel* |
| BioSemi | 70% | 70% | 70% | 80% |
| Smarting | 77.7% | 100% | 89% | 44% |
| DSI-24 | 82% | 64% | 72% | 55% |
| EPOC X | 71% | 42% | 0 | 57% |
| Mean across systems | 73% | 70% | 62% | 60% |

**Table 6:** Spatial-clustering results. Percentage of participants exhibiting a significant spatial cluster (p < 0.05 cluster corrected) corresponding to the N100 (negative clusters) and P300 (positive clusters) components in response to each stimulus, separately for each system.

| **System** | **N100** |  |  | **P300** |
| --- | --- | --- | --- | --- |
|  | *Standard* | *Target* | *Novel* | *Standard* |
| BioSemi | 100% | 80% | 70% | 90% |
| Smarting | 89% | 100% | 100% | 89% |
| DSI-24 | 81% | 72% | 81% | 72% |
| EPOC X | 28% | 28% | 14% | 42% |
| Mean across systems | 78% | 72% | 70% | 76% |

##### BioSemi

**Figure 5:**
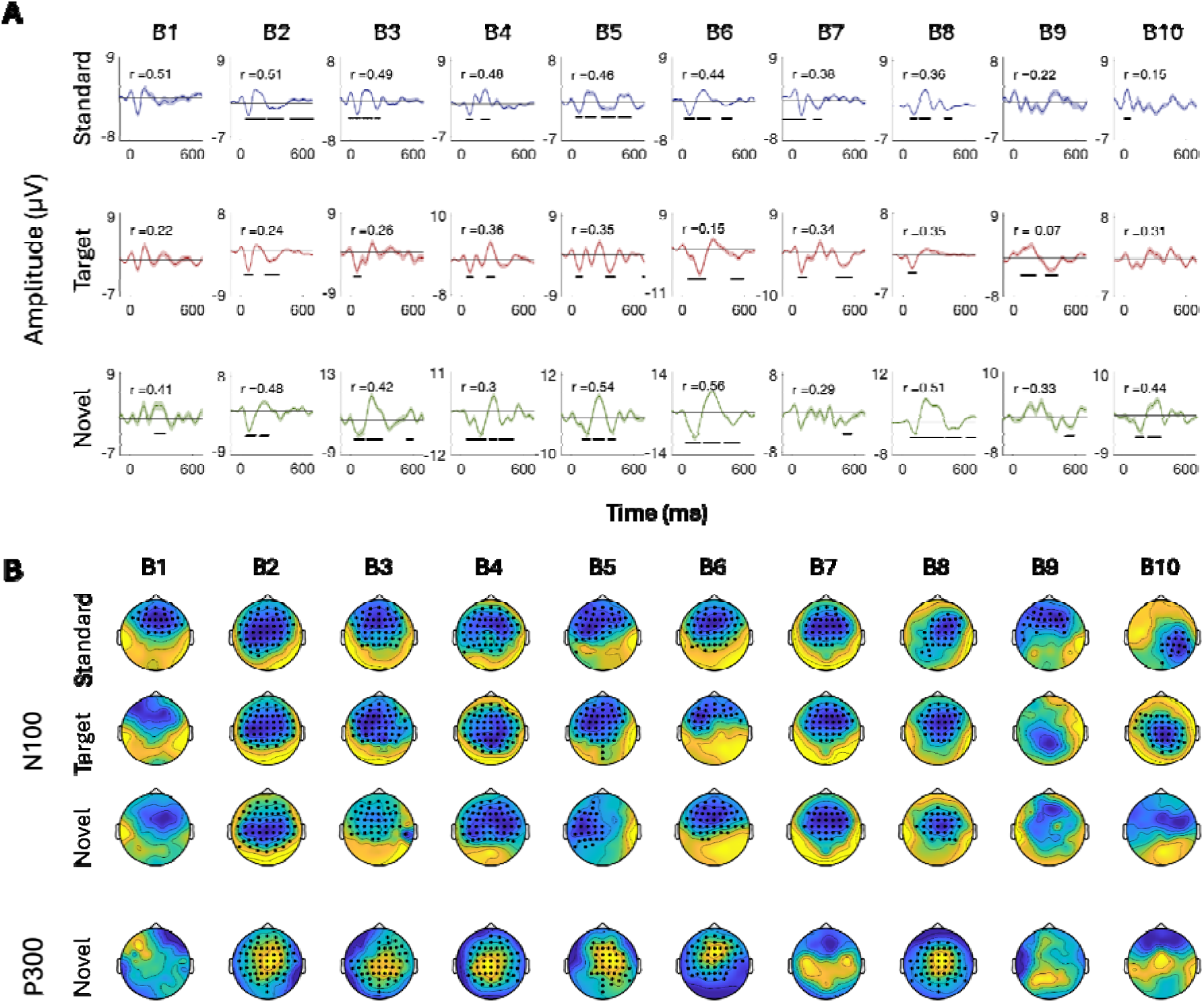
ERPs for all participants recorded with the BioSemi system. (A) ERPs in response to the three stimuli: Standard, Target, and Novel sounds, shown from electrode ‘Fz’. Shaded areas around the curves represent the standard error of the mean across trials. Time t=0 represents our best estimation of the onset of the stimulus, as determined based on direct audio-recordings synchronized to the EEG recordings via LSL (see Methods). Horizontal black lines indicate the time points in response to each stimulus where the ERP differs significantly from zero (p < 0.05, temporal-cluster corrected). Participants (B1-B10) are ordered in descending order based on their typicality score to Standard tones, which is represented by the r-value in each panel index. (B) Topographical maps showing the scalp distribution of the N100 (top) and P300 (bottom) responses, in a time-window ±10 ms around the group-level peak latencies. Black dots indicate electrodes where the response differed significantly from zero (one-sided p < 0.05, spatial-cluster corrected).

##### Smarting

**Figure 6:**
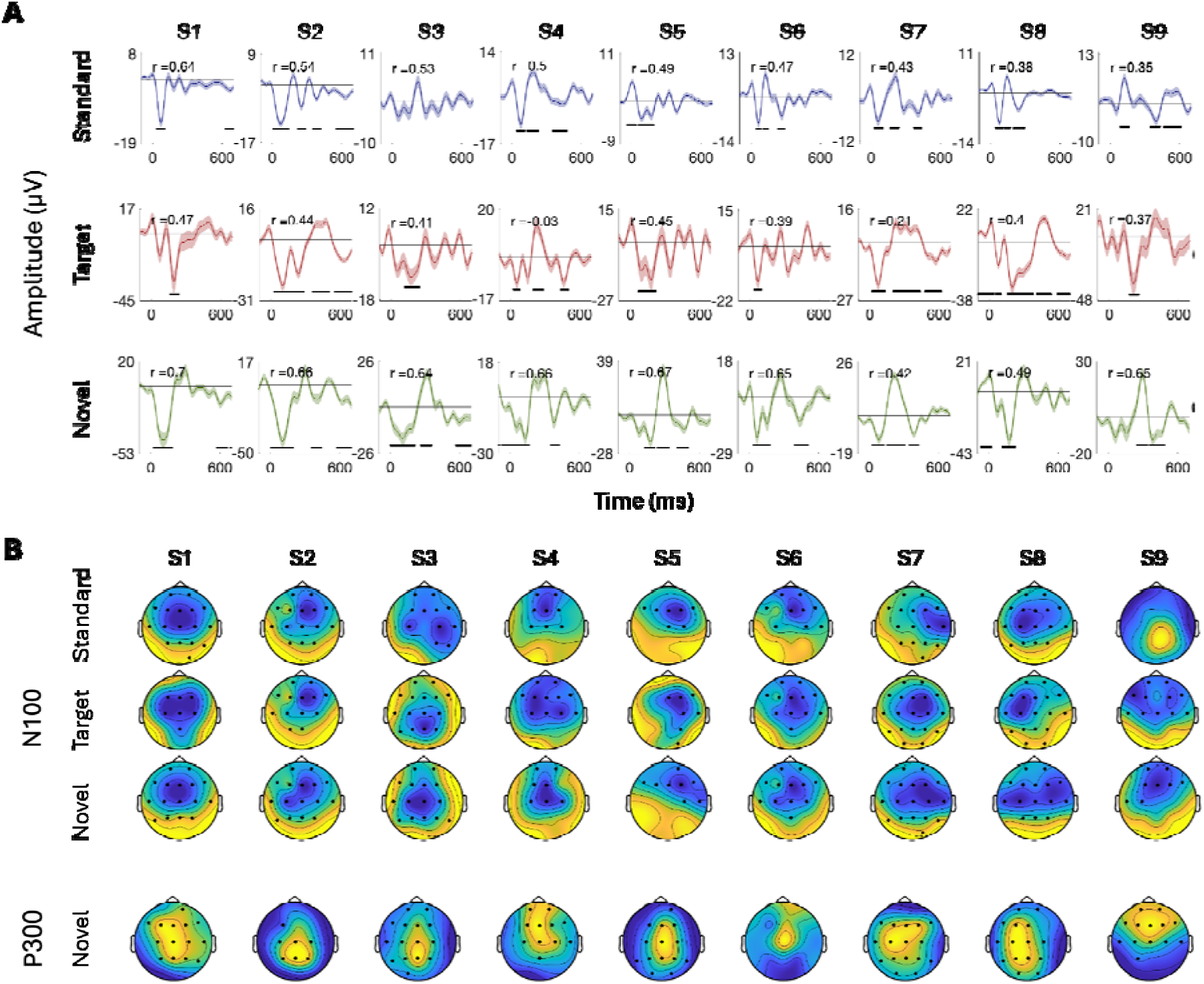
ERPs for all participants recorded with the in Smarting system. (A) ERPs in response to the three stimuli: Standard, Target, and Novel sounds, shown from electrode ‘Fz’. Shaded areas around the curves represent th standard error of the mean across trials. Time t=0 represents our best estimation of the onset of the stimulus, a determined based on direct audio-recordings synchronized to the EEG recordings via LSL (see Methods). Horizontal black lines indicate the time points in response to each stimulus where the ERP differs significantly from zero (p < 0.05, temporal-cluster corrected). Participants (S1-S9) are ordered in descending order based on their typicality score to Standard tones, which is represented by the r-value in each panel index. (B) Topographical maps showing th scalp distribution of the N100 (top) and P300 (bottom) responses, in a time window ±10 ms around the group-level peak latencies. Black dots indicate electrodes where the response differed significantly from zero (one-sided p < 0.05, spatial-cluster corrected).

##### DSI-24

**Figure 7:**
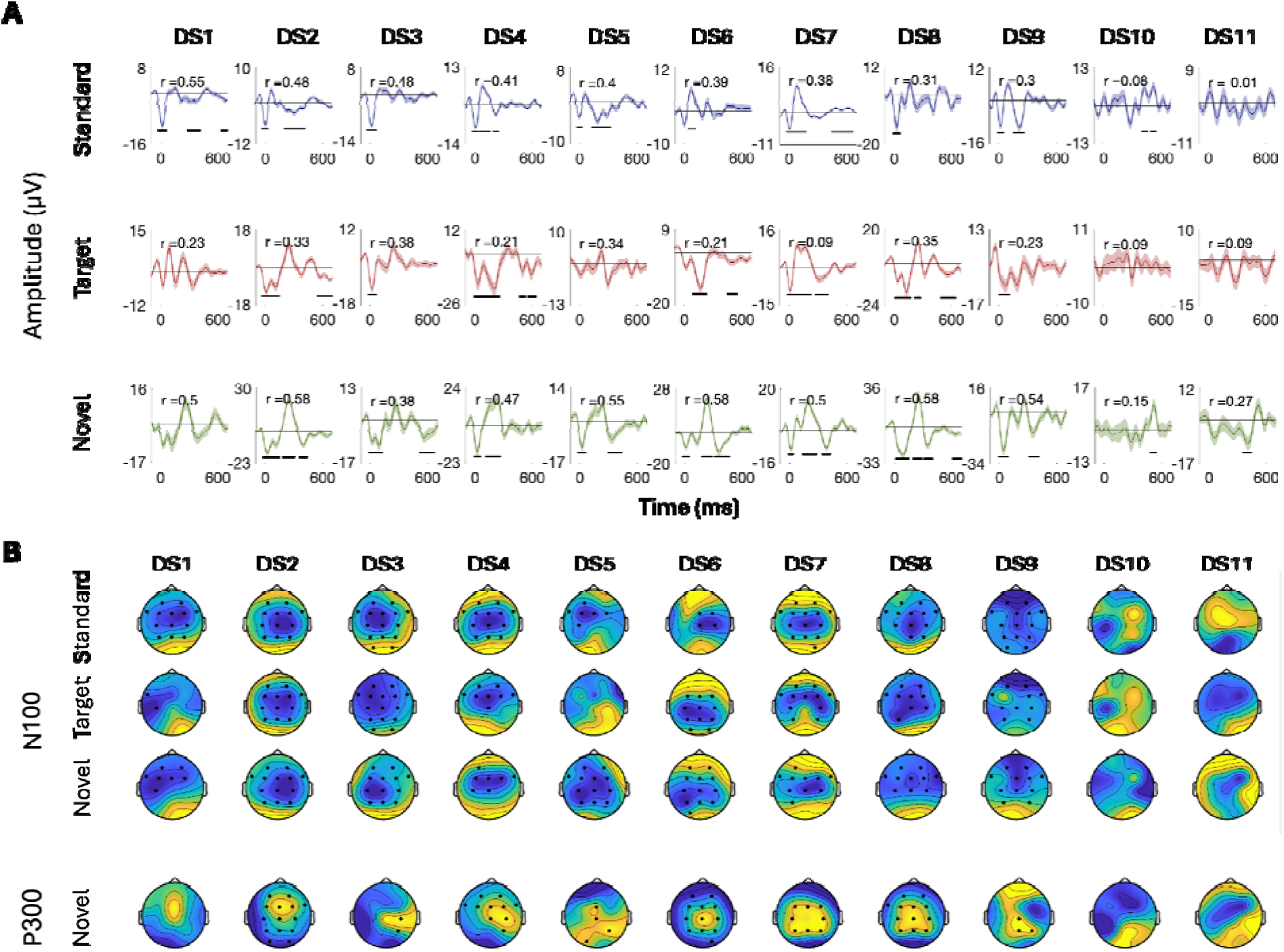
ERPs for all participants recorded with the DSI-24 system. (A) ERPs in response to the three stimuli: Standard, Target, and Novel sounds, shown from electrode ‘Fz’. Shaded areas around the curves represent the standard error of the mean across trials. Time t=0 represents our best estimation of the onset of the stimulus, as determined based on direct audio-recordings synchronized to the EEG recordings via LSL (see Methods). Horizontal black lines indicate the time points in response to each stimulus where the ERP differs significantly from zero (p < 0.05, temporal-cluster corrected). Participants (DS1-DS11) are ordered in descending order based on their typicality score to Standard tones, which is represented by the r-value in each panel index. (B) Topographical maps showing the scalp distribution of the N100 (top) and P300 (bottom) responses, in a time-window ±10 ms around the group-level peak latencies. Black dots indicate electrodes where the response differed significantly from zero (one-sided p < 0.05, spatial-cluster corrected).

##### EPOC X

**Figure 8:**
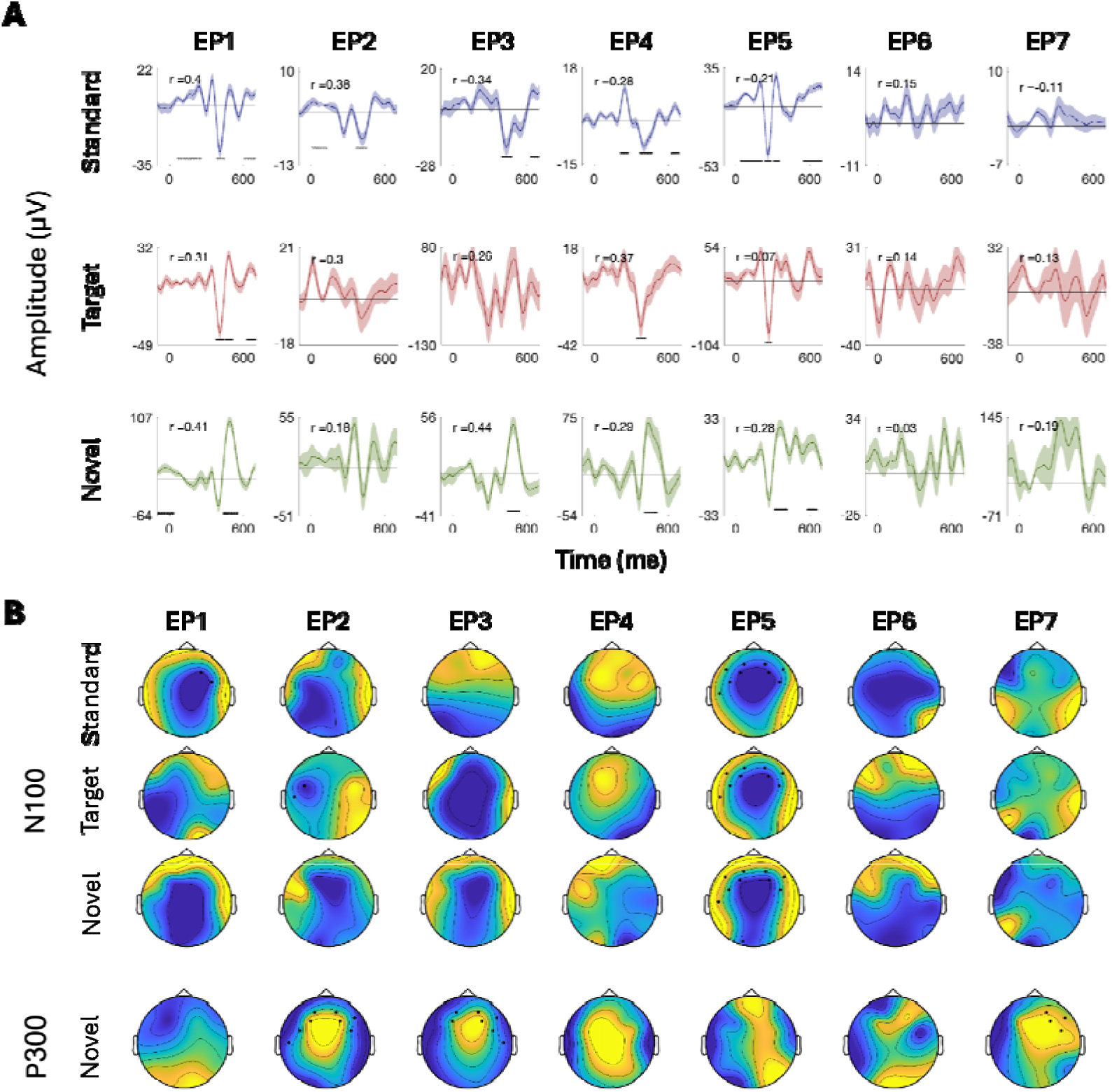
ERPs for all participants recorded with the EPOC-X system. (A) ERPs in response to the three stimuli: Standard, Target, and Novel sounds, shown from electrode ‘F4’. Shaded areas around the curves represent the standard error of the mean across trials. Time t=0 represents our best estimation of the onset of the stimulus, as determined based on direct audio-recordings synchronized to the EEG recordings via LSL (see Methods). Horizontal black lines indicate the time points in response to each stimulus where the ERP differs significantly from zero (p < 0.05, temporal-cluster corrected). Participants (EP1-EP7) are ordered in descending order based on their typicality score to Standard tones, which is represented by the r-value in each panel index. (B) Topographical maps showing the scalp distribution of the N100 (top) and P300 (bottom) responses, in a time-window ±10 ms around the group-level peak latencies. Black dots indicate electrodes where the response differed significantly from zero (one-sided p < 0.05, spatial-cluster corrected).

Last, to quantify the consistency in the time-course of responses across participants, we calculated each participant’s typicality score, a metric that reflects the correlation between that participant and all others (tested with the same system). The ERPs of individual participants in Figures 5-8 are ordered in descending order of typicality (based on the response to Standard tones) and typicality scores for all participants and stimuli are summarized in Figure 9, per system.

**Figure 9:**
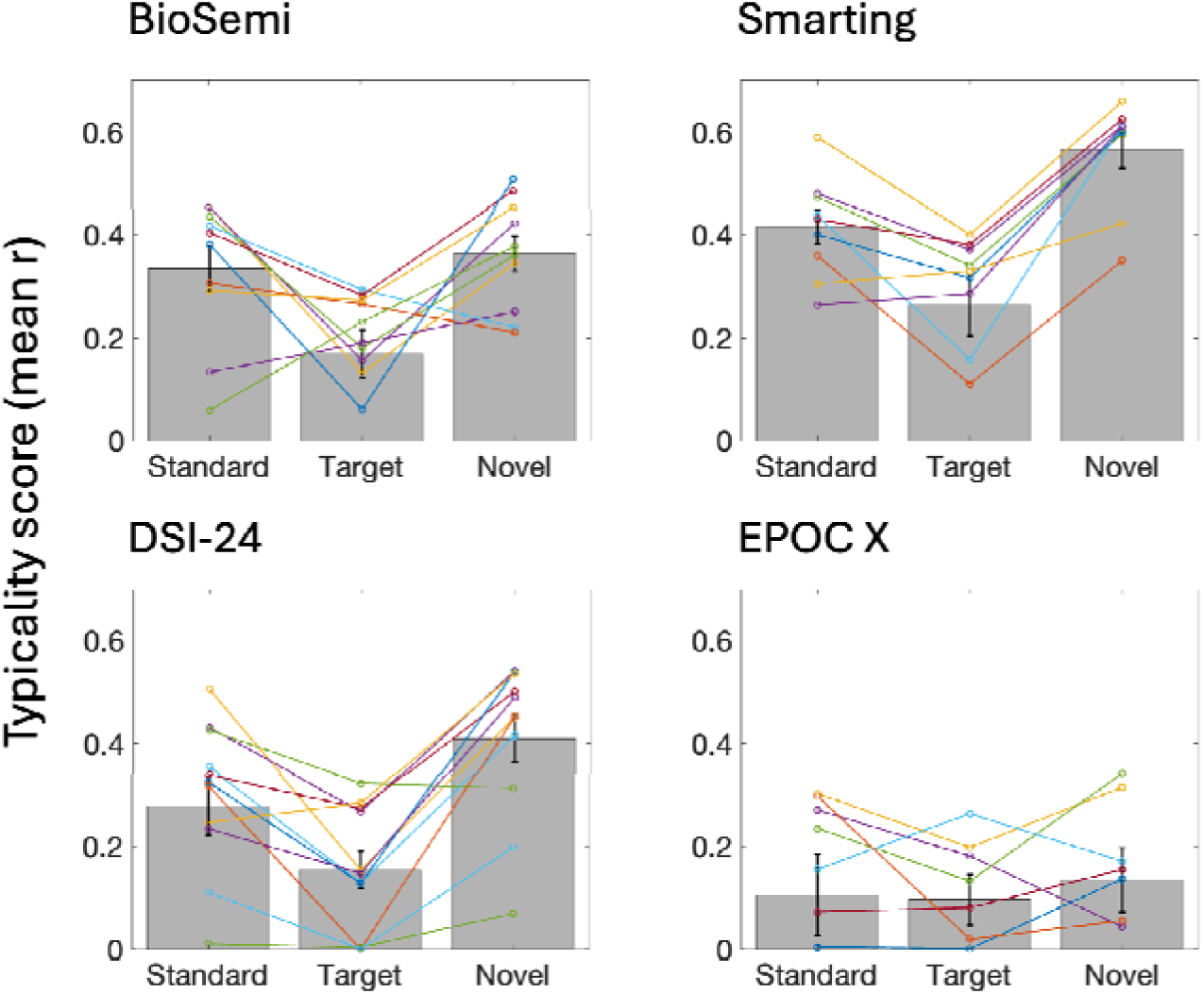
Typicality scores for each EEG system and stimulus. Bar graphs represent the mean typicality score in each condition (across participants tested with a given system), and error bars represent SEM. Colored lines connect the typicality scores of individual participants for the three stimuli.

A 3x4 repeated measures ANOVA revealed main effects of Stimulus (Standard, Target, Novel) and of the EEG system on typicality scores [Stimulus: F_(2,66)_ = 28.863, p < .001; System: F_(3,33)_=9.27, p <.001]. Post hoc pairwise comparisons (all bonferroni-corrected) showed that responses to Target tones had lower typicality scores than responses to Standard tones (t_(1,36)_ = 4.575, p < .001) and Novel sounds (t_(1,36)_ = 7.541, p < .001). In addition, responses to Novel sounds had higher typicality scores than responses to Standards (t_(1,36)_ = 2.965, p < .05). Regarding differences between EEG systems, the EPOC X system had significantly lower typicality scores relative compared to the other three systems (vs. BioSemi: t_(2,15)_ = 2.949, p < .05; vs. Smarting: t_(2,14)_ =5.273, p < .001; vs. DSI-24: t_(2,16)_ =3, p < .05). No other differences in typicality scores were found between EEG systems. All *p*-values were corrected for multiple comparisons using the Bonferroni correction.

#### Experiment 2 - Methods

##### Participants

EEG was recorded from 38 participants (30 females and 8 males), aged between 20 and 32 (M = 23.1±2.65). All participants were right-handed and reported having no hearing problems or diagnosis of any neurological or psychiatric condition. The study was approved by the IRB of Bar-Ilan University. Informed written consent was obtained from all participants. All participants were compensated for their participation with money or course credits.

#### Experiment setup and design

Participants performed the auditory oddball task used in Experiment 1 twice, in a within-session test-retest design. Between the two runs, participants listened passively to a 5–8 minute segment of natural speech – the results of which are the scope of the current study and are reported separately (Agmon et al., 2023).

EEG was recorded using the BioSemi Active II system with 64 channels, using a setup identical to that described in Experiment 1. Data were collected in an electrically shielded and acoustically attenuated room, and participants sat comfortably in front of a computer screen displaying visual instructions and a central fixation cross. Auditory stimuli were presented using headphones in a diotic presentation.

### Data Analysis

#### Behavioral Analysis

Analysis of behavioral responses followed the same procedure as in Experiment 1. A key press was considered a “hit” if it occurred within 200 to 1500 milliseconds after the target tone onset. Otherwise, the target was classified as a “miss.” All other key presses were categorized as “false alarms,” indicating responses that were either mistaken or unrelated to the stimuli.

#### EEG Analysis

##### Preprocessing

Preprocessing procedures were identical to those in Experiment 1. EEG data were re-referenced to linked mastoids, bandpass filtered (0.5–40 Hz), and detrended to retain the frequency range relevant to auditory ERPs. Independent component analysis (ICA) was applied to remove ocular and cardiac artifacts. Epochs were extracted from -100 to 400 ms relative to stimulus onset, and artifact-contaminated trials were excluded based on standard deviation thresholds.

##### Group-Level Analysis

For group-level analysis, grand averages were computed by averaging individual ERPs for each condition separately across participants. As we did in Experiment 1, after preprocessing, epochs were averaged separately for each condition (Standard, Target, Novel). Average signals were then baseline-corrected to the pre-stimulus period and low-pass filtered at 12 Hz (4th order zero-phase Butterworth filter), to produce ERPs. Grand average responses were derived by averaging the ERPs across participants, separately for each system and stimulus type.

We then used a data-driven clustering approach for each task condition to identify the time windows where the ERP significantly deviated from zero. For each task condition, a one-sample t-test was performed at each time point to identify periods where the response differed significantly from zero (p < 0.05). Next, a temporal cluster-based permutation test (Maris & Oostenveld, 2007) was conducted on the averaged neural signal from the Fz channel to identify significant time windows where ERPs deviated from zero (p < .05, cluster corrected). Early negative peak was classified as N100, and late positive as P300 peak based on waveform shape and topography.

##### Individual-Level Analysis

Individual-level analysis focused on the typicality score, described in Experiment 1, which quantifies the similarity of ERP waveforms across individuals. In each run, we calculated each participant’s typicality score by computing the average of all pairwise Pearson’s correlations between each participant’s ERP (at electrode Fz), and all other participants. This score represents how typical the response of each participant is compared to the group.

We then performed a 2×3 repeated-measures ANOVA with Run (first vs. second oddball run) and Stimulus (Standard, Target, Novel) as within-subjects factors. Additionally, we assessed the within-participant reliability of the typicality score between the two oddball runs, using Cronbach’s alpha test.

### Experiment 2 - Results

The grand-average ERPs elicited in response to Standard, Target, and Novel sounds, in each session, are shown in Figure 10. Similar to Experiment 1, temporal cluster-based analysis on the signal extracted from Fz reveals in both runs an early frontocentral negative peak around 90 ms, corresponding to the N100 component, followed by a wide frontocentral positive peak around 276 ms, corresponding to the P300 component. While the negative early pick was significant for all three conditions, the late positive peak was significant only for the Novel condition (p<0.05, cluster corrected).

**Table 7:** Time windows (in milliseconds) showing significant clusters (p < .05, cluster-corrected) corresponding to the N100 and P300 components in response to each task condition, presented separately for each run. Values in parentheses indicate the peak latency identified through visual inspection.

|  | **N100** |  |  | **P300** |
| --- | --- | --- | --- | --- |
|  | *Standard* | *Target* | *Novel* | *Novel* |
| First run | 62.5 – 133.7 (98) | 57.6 – 141.6 (90) | 65.4 – 133.8 (91) | 251.9 – 369.14 (315) |
| Second run | 63.4 – 118.1 (87) | 55.66 – 127.9 (90) | 63.4 – 124.4 (88) | 195.3 – 363.2 (276) |

**Figure 10:**
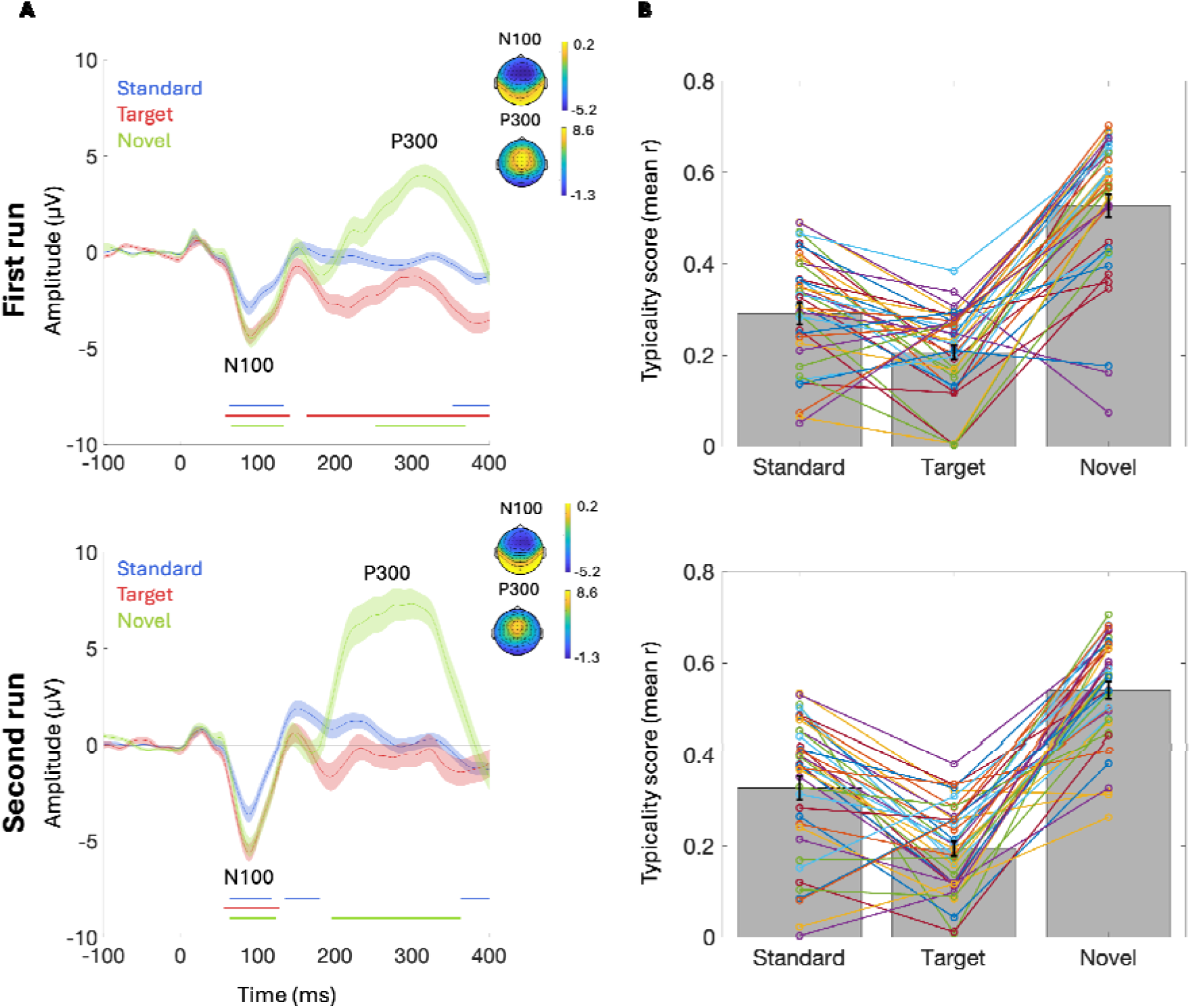
Results for Experiment 2 using BioSemi. (A) Grand average ERPs elicited in response to the three stimuli: Standard tones, Target tones, and Novel sounds, in the first (top) and second (bottom) oddball runs. ERPs are shown from electrode Fz. Shaded areas around the curves represent the SEM. Colored horizontal lines beneath each figur indicate the time-windows where the ERP in each condition differs significantly from zero (p < 0.05, cluster corrected). (B) The typicality scores in response to each stimulus, in the first (top) and second (bottom) oddball runs. Bar graphs represent the mean typicality score in each condition (across participants), and error bars represent SEM. Colored lines connect the typicality scores of individual participants for the three stimuli.

Analysis of ERP typicality scores for response to the three Stimuli replicates the results found in Experiment 1. There was a main effect of Stimulus [F_(2,158)_ = 156.4, p <.001], and post hoc contrasts revealed that typicality scores of the response to Target tones were lower compared to both Standard tones (t_(1,37)_ = 5.67, p < .001) and Novel sounds (t_(1,37)_ = 17.34, p < .001). Additionally, typicality scores were significantly higher in response to Novel vs. Standard sounds (t_(1,37)_ = 11.68, p < .001). Importantly, there was no main effect of Run, indicating that responses did not significantly differ across the two repetitions of the oddball task. Further supporting this, Cronbach’s Alpha test revealed high internal consistency of the typicality scores across the two oddball runs, for all stimuli (Standard = 0.93; Target = 0.78; Novel = 0.88). All *p*-values were adjusted for multiple comparisons across time windows using the Bonferroni correction.

## Discussion

In recent years, the landscape of EEG research has expanded dramatically with the advent of portable, user-friendly, and increasingly affordable mobile EEG systems. These advancements hold great promise for the use of EEG to identify biomarkers for cognitive, affective, and clinical traits at the individual level (Lau-Zhu et al., 2019). However, utilizing EEG-based markers as personalized neural metrics must be guided by quantification of their consistency and reliability within and between individuals. The current results contribute to these efforts, focusing on two of the most classic ERP components - the N100 and P300 and assessing their consistency across individuals and across four different EEG systems.

### Group level results

Results at the group-level replicate established patterns from prior auditory oddball studies showing an early N100 response to all auditory stimuli and a robust P300 component elicited mainly by the novel sounds (Intriligator & Polich, 1995; O’Connor et al., 1994; Segalowitz & Barnes, 1993; Tomé et al., 2015). The consistent emergence of the N100 and P300 components aligns with one of the foundational principles of EEG research: averaging across individuals produces strong and reliable neural responses, despite large differences in individual responses, anatomical variance, or external noise (Luck, 2014; Luck & Gaspelin, 2017). The finding that results were highly similar across three out of the four EEG systems is further reassuring and validates the signal quality of the two research-grade mobile EEG systems tested – the Smarting and DSI-24 – despite the different hardware, software, and experimental context under which data was collected (Gramann, 2024; Lau-Zhu et al., 2019; Mathewson et al., 2024, 2024). That said, the EMOTIV EPOC X device exhibited noisier ERP waveforms and greater temporal delays compared to all other systems, to the point that even group-level averages were barely reliable, let alone individual-level ERPs. This may partially be due to the different context under which the EPOC X data were collected (in-school EEG lab, with teenage students as participants), however it is unlikely that these factors fully explain the large discrepancy in results. A detailed investigation of the specific technical factors underlying the poorer signal quality of the EMOTIV is beyond the scope of this paper, but is in line with previous studies that also reported lower signal quality, reduced effect sizes, and higher proportion of artifacts in this system relative to research-grade systems (D’Angiulli et al., 2022; Duvinage et al., 2012, 2013), but see also (Badcock et al., 2013, 2015; Barham et al., 2017), These limitations are important to consider since the EMOTIV is by far the most affordable system, and therefore a prime choice for mobile-EEG studies (Sabio et al., 2024). Bearing this in mind, the group-level results indicate clearly that for research focusing on group-level averages, all four EEG systems are suitable for capturing canonical auditory ERP components. We next turn to discuss how these responses manifest at the individual level.

### Individual-level results

The question of how reliable individual-level ERP responses are, and how they relate to group-averaged signals, has been a central focus in ERP research for decades (Fabiani et al., 1998; Hajcak et al., 2017; Intriligator & Polich, 1995; Polich, 1987, 1997; Sandman & Patterson, 2000; Sklare & Lynn, 1984; Walhovd & Fjell, 2002). When focusing on the time windows and electrodes identified for the N100 and P300 in the group averages, we found that these components could be identified in most participants for the three systems that showed significant group-level responses (BioSemi, Smarting, and DSI-24), with 70–85% of participants showing statistically significant effects depending on the stimulus and system. For the EMOTIV system, detection rates were lower, reaching up to 60% depending on the stimulus. While this result is encouraging for precision imaging purposes, none of the stimuli or EEG systems produced responses that could be detected in *<u>all</u>*participants, which would be necessary for fully reliable individual-level neural metrics. This result aligns well with findings reported by Melnik and colleagues (2017), who found that individual differences accounted for 32% of the observed variance in ERP responses, across different participants, EEG systems, and tasks.

Interestingly, there were systematic differences in the detectability and typicality of the responses to the three different stimulus types used here: Novel sounds yielded a higher individual-level detection rate of the N100 and P300 responses, and ERP waveforms were more similar across individuals (high typicality) relative to Target and Standard sounds. This pattern was robust across all four EEG systems tested (although it was least reliable for the EPOC X), and was largely replicated across both experiments. Moreover, the finding in *Experiment 2* showing high test–retest reliability of responses to all three stimuli emphasizes the within-subject stability of individual-ERP waveforms despite substantial between-subject variations in their spatio-temporal patterns. Taken together, these results point to the superiority of ERPs to Novel sounds (in this experimental design) to serve as potential individual-level neural markers, at least in terms of individual-level sensitivity and waveform typicality, whereas responses to Target stimuli should be considered the least reliable. These notable differences between ERPs to Target and Novel sounds and their respective typicality and individual-level reliability resonate with discussions of underlying processes reflected by these ERPs, particularly the later components (Comerchero & Polich, 1999; Masson & Bidet-Caulet, 2019; Picton, 1992; Polich, 2007; Squires, et al., 1976). Specifically, while Novel sounds elicit a large P300 response (also referred to as the P3a), which is often attributed to an obligatory, near-automatic orienting response that is related to their unexpected nature (Debener et al., 2002, 2005; Parmentier & Andrés, 2010; Strobel et al., 2008), Target sounds are less surprising, but are expected to elicit late-responses associated with feature-based attention and detection, sometimes referred to as the P3b (Harmony et al., 2000; Näätänen, 2018; Picton, 1992; Schubert et al., 1998; Wickens, 1991). However, in the current data, there is little evidence for this P3b response in the group-level averages, across EEG systems and both experiments, and it is observed only in some of the individual time courses. This pattern suggests that while surprise-related responses are relatively ubiquitous across individuals, task-related neural responses and the detection of ‘to-be-attended’ stimuli are more variable across individuals, perhaps reflecting different strategies or attitudes towards the task (Peña & Cadaveira, 2000; Polich, 1987; Ritter et al., 1968; Simpson et al., 2014; Zeba et al., 2019). This result also resonates with findings by Segalowitz et al. (1993) who reported that the P2 and P3 responses to targets in a two-tone oddball paradigm were varied more substantially across individuals, potentially linked to inter-personal variation, relative to earlier components that were more similar across individuals. As a consequence, from a precision-imaging perspective, relying on responses to Targets as personalized neural metrics might be less optimal, as they are confounded by this inherent underlying variability. It is further important to note that Target-responses have actually often been proposed as useful markers for explaining individual differences, using both ‘simpler’ versions of the oddball paradigm (two-tone version; with only Standard and Target sounds (Fabiani et al., 1987; Melnik et al., 2017; Polich, 1987; Segalowitz & Barnes, 1993; Walhovd & Fjell, 2002), as well as more complex versions (e.g., P300 speller; Choi et al., 2023; Furdea et al., 2009; Halder et al., 2016; Käthner et al., 2013; Markovinović et al., 2022). Therefore, the utility of specific stimuli as individual markers may also vary as a function of the larger experimental context (Segalowitz & Barnes, 1993).

Given the potential promise of using EEG for precision imaging, it is imperative to distill which metrics could work well as potential individualized neural markers and which are inherently variable. Moving forward, these efforts could benefit from large open-source ERP datasets (for example, ERP CORE; Kappenman et al., 2021), or large-scale cross-site collaboration (e.g, the EEGManyLabs initiative; https://eegmanylabs.org) to refine methodological approaches and obtain reliable norms for individual-level ERPs. Such resources would enable researchers to better characterize baseline variability and more accurately identify individuals or cohorts that diverge from the norm, as well as track longitudinal and developmental trajectories. The current findings demonstrate the feasibility of using mobile EEG for this feat, with results largely generalizing across systems and contexts (with one exception), alongside emphasizing the importance of clear analytic strategies for quantifying inter-individual variability.

